# Origins of monkeypox viruses and their strains circulating in 2022

**DOI:** 10.1101/2022.09.07.506685

**Authors:** Ji-Ming Chen, Huan-Yu Gong, Rui-Xu Chen, Ming-Hui Sun, Yu-Fei Ji, Guo-Hui Li, Su-Mei Tan, Ying-Xue Sun

## Abstract

Monkeypox has spread unprecedentedly to nearly 100 countries and infected more than 51,000 people since 1 May 2022. This large-scale outbreak constituted a public health emergency of international concern (PHEIC), as declared by the World Health Organization. To better recognize and control this outbreak, we explore here through phylogenetic analysis the origins of MPXVs and their strains circulating in 2022, which remain unclear so far. Our results suggest that MPXVs possibly originated from some cowpox viruses and three lineages of MPXVs within Clade IIb circulated in 2022. Our results also suggest that two MPXVs respectively similar to the two MPXVs exported from Nigeria to the USA in 2021, ON676708/USA/2021 and ON676707/USA/2021, evolved into two lineages and sparked the large-scale outbreak in 2022, after their unknown evolutionary and epidemiological journeys possibly in Nigeria, the USA, or other countries before May 2022. This view does not stigmatize any country, because monkeypox is an endemic zoonosis in west Africa with wildlife reservoirs, and its large-scale outbreak in 2022 also resulted from the global decline of population immunity against smallpox due to the cease of vaccination against smallpox decades ago.

## Introduction

To date, monkeypox, which is a zoonotic endemic disease in regions of Africa caused by monkeypox virus (MPXV), has spread unprecedentedly to nearly 100 countries with more than 51,000 confirmed human cases and 15 confirmed deaths since 1 May 2022 (1-4). This large-scale outbreak, which was abbreviated below as MPX/outbreak/2022, constituted a public health emergency of international concern (PHEIC), as declared by the World Health Organization (WHO) (1). Many countries have strengthened a series of measures to stamp out the outbreak.

MPXV virions are enveloped with brick-shaped geometries around 200 nm wide and 250 nm long. The genome of MPXV is a linear double-stranded DNA (≈197 kb) which is more conserved in the central regions than in the terminal regions. MPXV constitutes the species *Monkeypox virus* and shares the same genus *Orthopoxvirus*, subfamily *Chordopoxvirinae* and family *Poxviridae* with *Camelpox virus* (CMLV), *Cowpox virus* (CPXV), *Raccoonpox virus, Skunkpox virus, Vaccinia virus* (VACV), *Variola virus* (VRAV, the etiology of smallpox), *Ectromelia virus* (ECTV) and four other species (3).

To better recognize and control MPX/outbreak/2022, we explore here through phylogenetic analysis the origins of MPXVs and their strains circulating in 2022, which remain unclear so far.

## Methods and Materials

### Virus designation

Virus strains were designated using their species names (or the relevant abbreviations) and GenBank or GISAID accession numbers followed by the countries and years where or when the strains circulated, or by strain or host names. The Democratic Republic of the Congo, United Kingdom, Unites States, and Central African Republic are abbreviated as DRC, UK, USA, and CAR. The species name MPXV was omitted in relevant designations.

### Phylogenetical analysis

Phylogenetical relationships were calculated using the software package MEGA (version X) with the method of neighbor-joining and the nucleotide substitution model of maximum composite likelihood (5). Gaps were treated as partial deletion (the cutoff value = 80%). The relationships were also calculated using the method of maximum likelihood and the nucleotide substitution model with the lowest Bayesian Information Criterion (BIC) score (the model was considered to describe the substitution pattern the best) (5). The results of these two methods both supported the opinions of this analysis.

## Results and Discussion

### Origin of CPXVs

We downloaded all genomic sequences (>100,000 bp) of orthopoxviruses available in GenBank excluding MPXVs isolated in 2022. If the sequence identities of two or more strains from the same country and the same host were less than 0.008%, some of these strains were excluded for the phylogenetic analysis. The analysis showed that each species of orthopoxviruses was located in a specific branch of the phylogenetic tree, except that CPXVs occupied several branches in the tree, which represented Lineages 1−3 CPXVs (Fig. 1A). This is consistent with a previous report (6). The topology of Fig. 1A suggested with solid support of bootstrap values (100%) that MPXVs and VACVs possibly originated from Lineage 3 CPXVs and that CMLVs and *Taterapox virus* possibly originated from Lineage 2 CPXVs. Meanwhile, Fig. 1A suggested that CMLVs and *Taterapox virus* can be considered as the ancestors of VARVs. Consistent with previous studies (7), Fig. 1A showed that humans, non-human primates, and various other mammals support CPXV infection, which could facilitate some CPXVs to evolve into new species. In phylogenetics, CPXVs were likely situated at a peak of a “hill”, and they evolved along different “downhill” routes into multiple new species. Fig. 1A is not fully novel as compared with previous studies (6,7), but its application here in exploring the origins of MPXV and other species is novel.

**Fig. 1.**
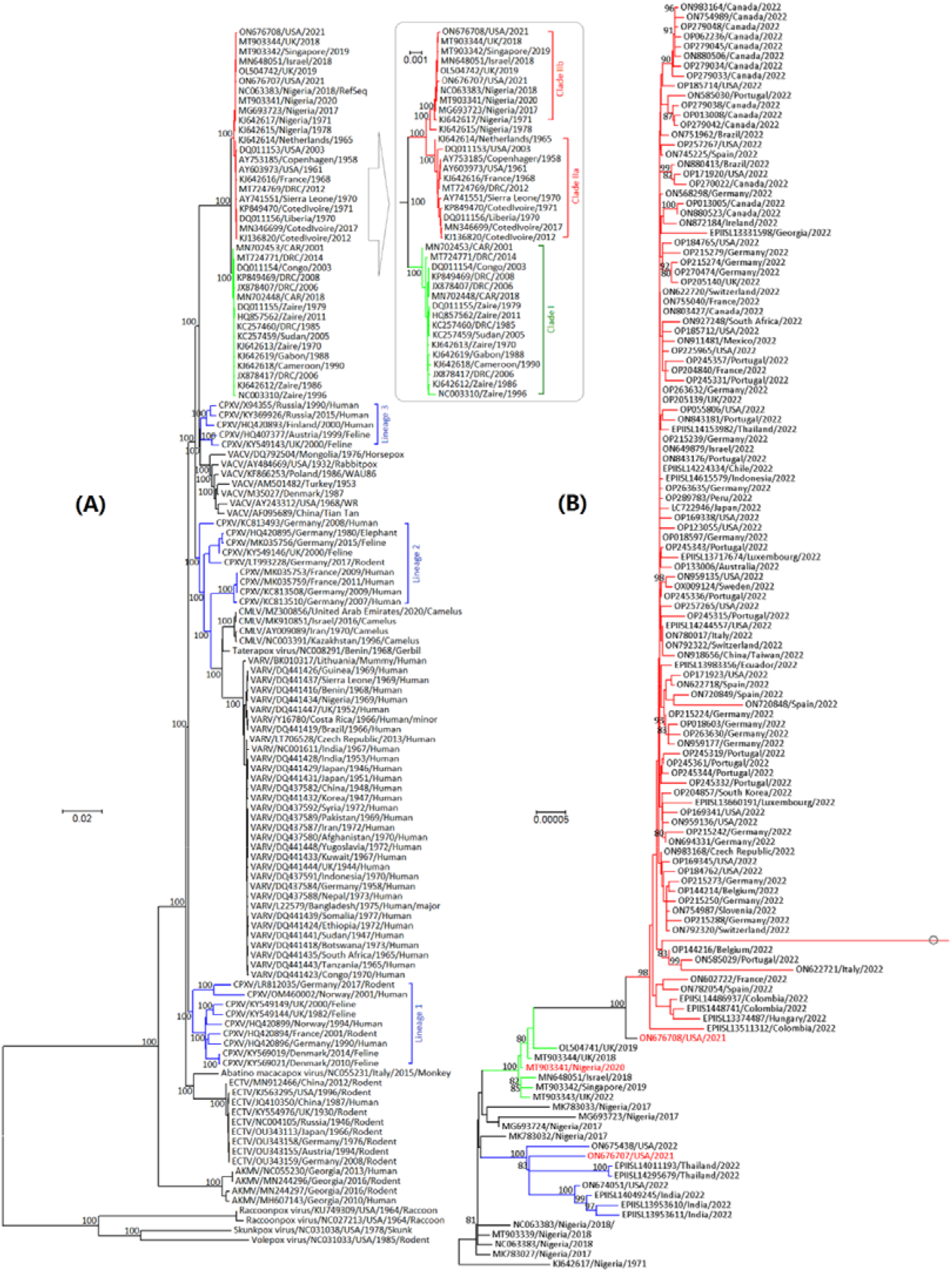
Phylogenetic relationships among orthopoxviruses **(A)** and Clade IIb monkeypox viruses (MPXVs) **(B). A:** the blue branches represent three lineages of cowpox viruses, and the green and red branches represent Clades I and II of MPXVs, which are shown separately nearby with an enlarged scale of genetic distances. **B:** the red, green, and blue branches represent Lineages A−C MPXVs; the branch with a circle represents the strain OP012849/China/2022; the three strains in red letters represent the ones circulating before 2022 the closest in phylogenetics to the three lineages of MPXVs circulating in 2022, respectively.

### Clades of MPXVs

Fig. 1A shows that MPXVs can be classified into Clades I and II, and Clade II includes Clades IIa and IIb, as per the novel nomenclature issued by the WHO (1). Consistent with previous studies (8,9), Fig. 1A further shows: Clade I mainly circulates in central Africa (CAR, DRC, Cameroon, Gabon, Congo, and Sudan); infections of Clade I MPXVs outside Africa through international transportation have not been reported; KJ642615/Nigeria/1978 was an intermediate strain between Clades IIa and IIb; Clade IIa mainly circulates in the western African countries of Sierra Leone, Liberia, and Cote d’Ivoire; infected humans or animals with Clade IIa MPXVs were transported to Denmark in 1958, the USA in 1961 and 2003, Netherlands in 1965, and France in 1968; Clade IIb has been circulating in the western African country of Nigeria at least after 1971; infected humans with Clade IIb MPXVs were transported to the UK in 2018 and 2019, Israel in 2018, Singapore in 2019, and the USA in 2021.

### Origins of the MPXVs circulating in 2022

We downloaded all genomic sequences (>100,000 bp) of MPXVs available in GenBank and GISAID. If sequence identities of two or more strains from the same country in the same year were less than 0.003%, some of them were excluded for phylogenetic analysis. The initial analysis showed that the MPXVs circulating in 2022 belonged to Clade IIb and hence Clade I MPXVs were excluded in the final analysis which was shown in Fig. 1B.

The MPXVs circulating in 2022 were located in three lineages within Clade IIb with strong support of bootstrap values (98−100%) (Fig. 1B). Lineage A, which has circulated in many countries in 2022, likely originated from a variant represented by the strain ON676708/USA/2021. ON676708/USA/2021 was transported from Nigeria to Maryland in November 2021 (9). Lineage B, which has circulated in 2022 in the countries of the USA, Thailand, and India (10), likely originated from a variant represented by the strain ON676707/USA/2021. ON676707/USA/2021 was transported from Nigeria to Texas in July 2021 (8,9). Lineage C, which circulated in the UK in 2022, likely circulated in Nigeria in 2017−2020 and was sporadically transported to Israel, Singapore, and the UK in 2018−2019.

These above data suggest that two MPXVs in 2021, which were respectively similar to the two MPXVs exported from Nigeria to the USA, ON676708/USA/2021 and ON676707/USA/2021, evolved into Lineages A and B and sparked MPX/outbreak/2022, after their unknown epidemiological and evolutionary journeys possibly in Nigeria, the USA, or other countries before May 2022. This view does not stigmatize any country, because monkeypox is an endemic zoonosis in west Africa with wildlife reservoirs, and MPX/outbreak/2022 also resulted from the global decline of population immunity against smallpox due to the cease of vaccination against smallpox decades ago (1,3,8,9).

## Conclusions

The analysis suggests that MPXVs possibly originated from some CPXVs and three lineages of Clade IIb MPXVs circulated in 2022. It also suggests that two MPXVs in 2021 respectively similar to the two MPXVs exported from Nigeria to the USA, ON676708/USA/2021 and ON676707/USA/2021, evolved into two lineages and sparked MPX/outbreak/2022, after their unknown epidemiological and evolutionary journeys possibly in Nigeria, the USA, or other countries before May 2022.

## Author contributions

J.M.C conceived, designed, and supported this study, analyzed the data, and drafted the manuscript. Y.X.S made the core conclusion, analyzed the data, and revised the manuscript. H.Y.G, R.X.C, S.M.H, Y.F.J, G.H.L, and T.S.M. collected and analyzed relevant data and revised the manuscript.

## Acknowledgments

This study was supported by the High-Level Talent Fund of Foshan University (no. 20210036). The funder did not play any role in this study.

